# Close proximity interactions support transmission of ESBL-*K. pneumoniae* but not ESBL-*E. coli* in healthcare settings

**DOI:** 10.1101/413500

**Authors:** Audrey Duval, Thomas Obadia, Pierre-Yves Boëlle, Eric Fleury, Jean-Louis Herrmann, Didier Guillemot, Laura Temime, Lulla Opatowski, the i-Bird Study group

## Abstract

Antibiotic-resistance of hospital-acquired infections is a major public health issue. The worldwide emergence and diffusion of extended-spectrum β-lactamase (ESBL)-producing Enterobacteriaceae, including Escherichia coli (ESBL-EC) and Klebsiella pneumoniae (ESBL-KP), is of particular concern. Preventing their nosocomial spread requires understanding their transmission. Using Close Proximity Interactions (CPIs), measured by wearable sensors, and weekly ESBL-EC– and ESBL-KP–carriage data, we traced their possible transmission paths among 329 patients in a 200-bed long-term care facility over 4 months. Based on phenotypically defined resistance profiles to 12 antibiotics, new bacterial acquisitions were tracked. Extending a previously proposed statistical method, the CPI network’s ability to support observed incident colonization episodes of ESBL-EC and ESBL-KP was tested. Finally, mathematical modeling based on our findings assessed the effect of several infection-control measures. A potential infector was identified in the CPI network for 80% (16/20) of ESBL-KP acquisition episodes. The lengths of CPI paths between ESBL-KP incident cases and their potential infectors were shorter than predicted by chance (P = 0.02), indicating that CPI-network relationships were consistent with dissemination. Potential ESBL-EC infectors were identified for 54% (19/35) of the acquisitions, with longer-than-expected lengths of CPI paths. These contrasting results yielded differing impacts of infection control scenarios, with contact reduction interventions proving less effective for ESBL-EC than for ESBL-KP. These results highlight the widely variable transmission patterns among ESBL-producing Enterobacteriaceae species CPI networks supported ESBL-KP, but not ESBL-EC spread. These outcomes could help design more specific surveillance and control strategies to prevent in-hospital Enterobacteriaceae dissemination.

**Author summary:** Tracing extended-spectrum β-lactamase (ESBL) dissemination in hospitals is an important step in the fight against the spread of multi-drug resistant bacteria. Indeed, understanding ESBL spreading dynamics will help identify efficient control interventions. In the i-Bird study, patients and hospital staff from a French long-term care facility in France carried a wearable sensor to capture their interactions at less than 1.5 meters, every 30 seconds over a 4-month period. Every week, patients were also swabbed to detect carriage of ESBL-producing Enterobacteriaceae. Based on the analysis of these longitudinal data, this study shows that ESBL-producing *Klebsiella pneumoniae* (ESBL-KP) mostly spreads during close-proximity interactions between individuals, while this is not the case for ESBL-producing *Escherichia coli* (ESBL-EC), suggesting that ESBL-KP but not ESBL-EC may be controlled by contact reduction interventions.

## Introduction

Multidrug resistant (MDR)-Enterobacteriaceae are a common cause of healthcare-associated and community-acquired infections in humans (1), due to the increase over recent years of third-generation cephalosporin, fluoroquinolone and carbapenem resistances (2,3), leading to difficulties finding appropriate treatment and increased mortality and morbidity. The recent emergence of colistin resistance among Gram-negative bacteria also raises new concerns (4). According to a recent World Health Organization (WHO) assessment, the greatest threats to human health is posed by extended-spectrum β-lactamase (ESBL)-producing Escherichia coli (ESBL-EC) and ESBL-producing Klebsiella pneumoniae (ESBL-KP) (5), causing bloodstream, urinary tract and respiratory infections mostly (3).

The infections burden of those bacteria are predominantly in hospitals worldwide. In a WHO review, E. coli (20.1%) was the most frequent single pathogen causing healthcare-associated infections in mixed patient populations (6). A large US prevalence survey found E. coli and K. pneumoniae to be responsible for 20% of all healthcare-associated infections and 50% of healthcare-associated urinary tract infections (7). In a recent pan-European cohort, bacteremia caused by ESBL-producing Enterobacteriaceae increased mortality (hazard ratio (HR): 1.63; 95% confidence interval (CI): 1.13 – 2.35), lengths of stay (4.9; 95% CI: 1.1 – 8.7 days), and healthcare-associated costs compared with non-ESBL-producing strains (8). However, to control the threat of these bacteria in hospital settings, more insight is needed regarding their transmission routes (9).

New technologies to measure close proximity interactions (CPIs) by wireless sensors (10,11) have been implemented in hospital investigations (12–16). CPIs are assumed to be a proxy of human contacts that support human-to-human microorganisms transmission (17–20). In an earlier study, CPI networks were shown to be a significant support of Staphylococcus aureus transmission (21,22).

In this study, we exploited the original longitudinal observational i-Bird (Individual-Based Investigation of Resistance Dissemination) data collected in a 200-bed long-term care facility (LTCF). CPIs between patients and hospital staff were recorded every 30 s over a 4-month period and rectal swabs were collected weekly from patients to test for Enterobacteriaceae carriage. We separately examined the role of CPIs in ESBL-EC and ESBL-KP spread (9). Using a mathematical model, we tested whether CPI information could be useful in designing and organizing control interventions in hospital.

## Results

### ESBL-EC and ESBL-KP colonization

The i-Bird study included 329 patients. The weekly average carriage prevalence of ESBL-producing Enterobacteriaceae was 16.8%. The predominant species were E. coli and K. pneumoniae, with on average 11.5% of patients colonized weekly by an ESBL-EC, and 3.7% by an ESBL-KP (Table 1). Over the 4 months of the study, 203 patients were admitted and swabbed at admission (Figure S3); 16 of those patients carried an ESBL-EC and 2 an ESBL-KP on admission, representing respective importation rates of 8% and 1%. Overall, 35 incident colonization episodes were observed for ESBL-EC (acquisition rate: 0.66%/week), and 20 for ESBL-KP (acquisition rate: 0.38%/week).

**Table 1.** Characteristic of the extended-spectrum β-lactamase ESBL-producing E. coli (ESBL-EC)- and K. pneumoniae (ESBL-KP)-carrier population. Details about colonized patients, ward prevalence, incidence and CPIs description of colonized patients are summarized below.

| Characteristic | Ward 1 | Ward 2 | Ward 3 | Ward 4 | Ward 5 | Hospital |
| --- | --- | --- | --- | --- | --- | --- |
| No. of patients per week (median) | 36 | 27 | 21 | 28 | 16 | 128 |
| ESBL-EC |  |  |  |  |  |  |
| Age (median (range)) | 53.23 (31–70) | 54.2 (40–70) | 57.5 (32–80) | 48.25 (27–80) | 84.36 (76–100) | 60.82 (27–100) |
| Gender (% female) | 53.85 | 40 | 62.5 | 50 | 63.64 | 33.33 |
| Total number of colonized patients by at least one ESBL-EC | 13 | 5 | 8 | 8 | 11 | 45 |
| Average weekly prevalence (%) | 12.79 | 8.38 | 8.41 | 7.66 | 14.85 | 11.51 |
| Average incidence (acquisitions/100 patients/week) | 2.71 | 1.16 | 1.35 | 1.45 | 4.23 | 1.96 |
| Mean no. of daily distinct CPIs (SD) | 16.33 (10.1) | 9.97 (2) | 12.71 (4) | 13.7 (4) | 7.87 (2.3) | 12.44 (6.7) |
| With patients | 8.11 (6.7) | 3.52 (1.5) | 6.56 (2.6) | 7.42 (3.9) | 3.11 (0.8) | 5.98 (4.5) |
| With hospital staff | 8.22 (3.7) | 6.45 (1.2) | 6.16 (2) | 6.29 (2.4) | 4.76 (1.8) | 6.47 (2.8) |
| Mean daily cumulative duration of CPI (SD) | 43.83 (13.9) | 32.42 (18.2) | 28.15 (26.7) | 27.62 (14.8) | 54.36 (26.7) | 39.47 (22.5) |
| With patients | 89.54 (44.4) | 57.11 (40.3) | 43.47 (36.4) | 46.93 (42.4) | 109.54 (49) | 75.06 (49.5) |
| With hospital staff | 8.05 (2.4) | 15.62 (13.7) | 15.71 (19.6) | 13.68 (9) | 15.24 (25) | 13.01 (15.7) |
| ESBL-KP |  |  |  |  |  |  |
| Age (median (range)) | 53 (34–70) | 40 | 53.5 (27–70) | 53 (44–62) | 0 | 52.39 (27–70) |
| Gender (% female) | 10 | 100 | 75 | 100 | 0 | 6.67 |
| Total number of colonized patients by at least one ESBL-KP | 10 | 1 | 4 | 3 | 0 | 18 |
| Average weekly prevalence (%) | 14.4 | 0.26 | 1.61 | 0.73 | 0 | 3.73 |
| Average incidence (acquisitions/100 patients/week) | 4.16 | 0 | 1.13 | 0.3 | 0 | 1.15 |
| Mean no. of daily distinct CPIs (SD) | 13.61 (1.2) | 12 | 13.56 (5.8) | 8.17 (0.8) | 0 | 12.6 (3.3) |
| With patients | 6.48 (1.4) | 5.96 | 7.99 (4.4) | 4.26 (0.4) | 0 | 6.42 (2.4) |
| With hospital staff | 7.13 (1.2) | 6.04 | 5.57 (3) | 3.9 (1.1) | 0 | 6.19 (2) |
| Mean daily cumulative duration (SD) | 52.33 (18.6) | 7.7 | 25.01 (26.3) | 33.22 (7.9) | 0 | 40.6 (22.9) |
| With patients | 105.52 (50.2) | 10.75 | 36.12 (36.6) | 47.21 (7.9) | 0 | 75.12 (53.5) |
| With hospital staff | 10.64 (3.4) | 3.85 | 13.34 (19.8) | 16.65 (8.2) | 0 | 11.86 (9.6) |

Prevalence and incidence of ESBL-EC were the highest in the geriatric ward 5, with a weekly average of 15% of colonized patients, and an incidence of 4% per week; whereas ESBL-KP prevalence and incidence were the highest in the neurology ward 1 (14% and 4% per week, Table 1). Colonized ward 1 patients had the highest average daily distinct CPIs (16.33 ± 10.1 CPIs per day and 13.61 ± 1.2 CPIs per day, for ESBL-EC and ESBL-KP respectively), equally distributed in patients and hospital staff, while the highest cumulative CPI duration was found for ESBL-EC–colonized ward 5 patients.

### Are ESBL-EC and ESBL-KP transmissions supported by CPIs?

Two ESBL-producing Enterobacteriaceae strains were assumed to be identical when they belonged to the same species (EC or KP) and had the same resistance-status pattern to 12 selected antibiotics, allowing for susceptible to intermediate (S–I) or intermediate to resistance (I–R) differences. Thirty-five incident-colonization episodes (in which a patient was found to be colonized during a given week by a strain he/she was not carrying the preceding week) were identified for ESBL-EC and 20 for ESBL-KP. For each incident-colonization episode, “transmission candidates”, i.e. patients who carried the same strain as the case over the preceding 4 weeks, were identified; among transmission candidates, those who were linked to the case via the shortest distance (defined as the number of edges between the two, i.e. length of CPI path) on the CPI-network were called “potential infectors”. For both species, incident-colonization episodes were mostly resolved during the preceding week: a potential infector had been identified during the first week preceding the episode for 56% and 63% of all ESBL-KP and ESBL-EC episodes respectively.

To determine whether CPIs could explain transmission, we tested whether observed distances along the CPI-network between a case and their closest potential infectors were comparable to distances expected under the null hypothesis of independence between CPIs and carriage data. Expected distances were computed as the average of distances obtained from 200 simulations using randomly permutated carriage data.

Transmission of ESBL-EC. No carrier of the same strain was found over the preceding 4 weeks (transmission candidate) for 13 of the 35 incident-colonization episodes. For three additional episodes, no potential infector was found throughout the network, resulting in a total of 16/35 unresolved episodes. Observed and expected case-to-potential infector distances did not differ significantly based on the remaining 19 resolved episodes (P = 0.25, Wilcoxon signed rank paired test). Indeed, more direct CPIs (i.e., distance-1) between cases and their closest potential infectors were found in the permutated data than in the observed data (Figure 1A, 20% and 5% respectively). Conversely, more distance-2 were found in the observed data.

**Figure 1.**
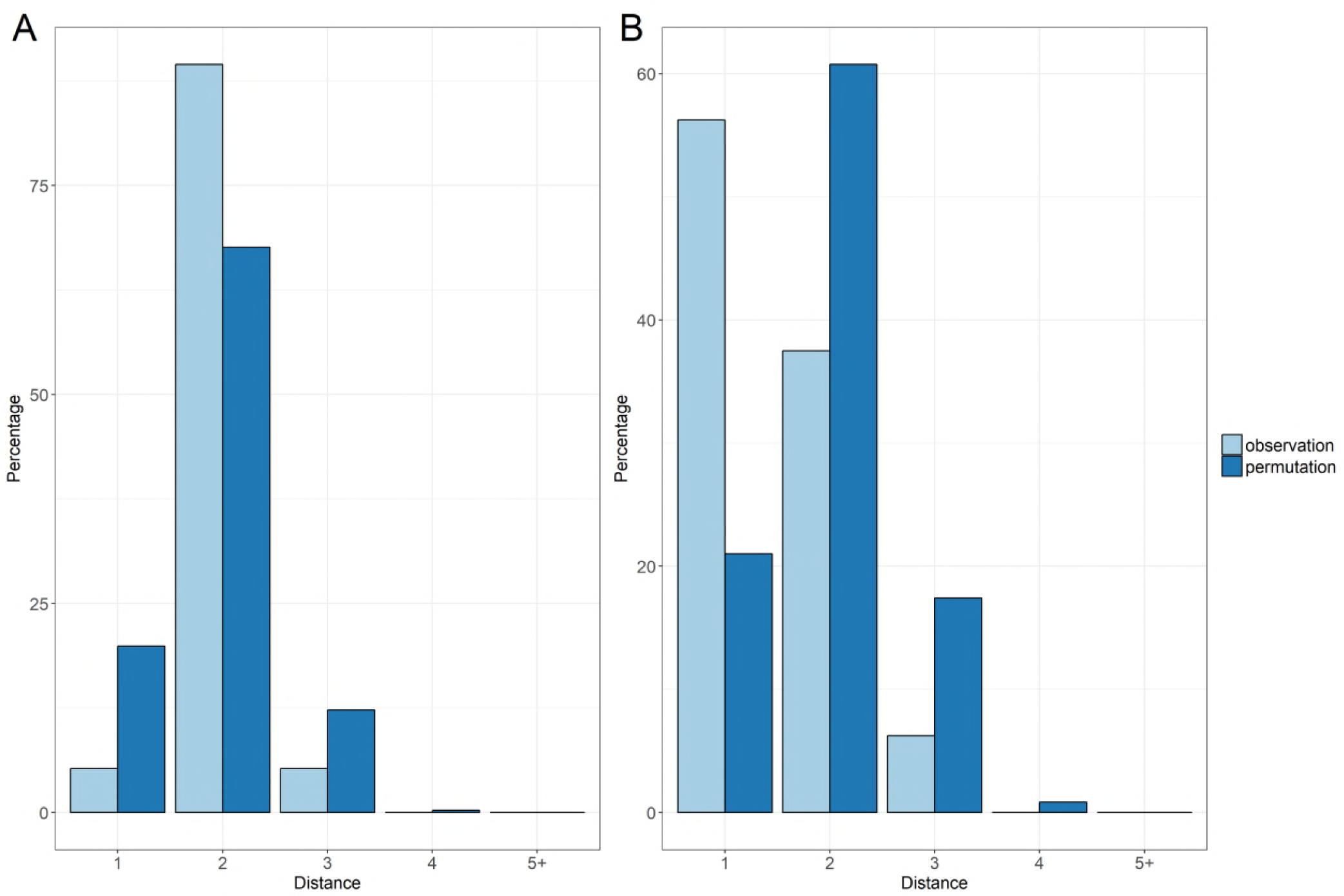
Distribution of distances between acquisition cases and their closest potential infector. Comparison between observed data (light blue) and random permutated data (dark blue). For each incident colonisation case, potential infectors were selected as the closest in the CPI-network of all candidates sharing the same strain as the case in the preceding 4 weeks. (A) ESBL-EC distribution. (B) ESBL-KP distribution.

Transmission of ESBL-KP. Only four of the 20 episodes were not resolved for ESBL-KP i.e. no potential infectors were found. The case-to-potential infector distances differed significantly between observed and simulated datasets for the 16 resolved episodes. That distance was shorter than expected by chance (P = 0.025, Wilcoxon signed rank paired test), suggesting that ESBL-KP transmission was indeed supported by CPIs. There were also more direct CPIs (distance-1) between incident-colonization episodes and their closest potential infector than expected by chance (Figure 1B, 56% vs. 21%).

Intermediaries. When looking more precisely at distance-2 between incident cases and their closest potential infector, and more particularly at the distribution of status of intermediaries (i.e. patients or hospital staff), observed and permutated data differed clearly for both species. More patient intermediaries in the observed data for ESBL-KP and more hospital staff for ESBL-EC (Figure S4) were found.

### Sensitivity analyses

Sensitivity of the results to strain definition. As expected, a stricter definition of identity between strains, taking into account S–I or I–R mismatches in addition to S–R, led to identifying more incident-colonization episodes (49 vs. 35 for ESBL-EC and 49 vs. 20 for ESBL-KP, Table 2). With this scenario, lower percentage of episodes were resolved (41% vs. 54% for ESBL-EC and 33% vs. 80% for ESBL-KP), especially for ESBL-KP but in absolute values, almost same numbers of episodes were resolved (20/49 vs. 19/35 for ESBL-EC. and 16/49 vs. 16/20 for ESBL-KP). Pertinently, application of the strict definition did not change previous conclusions. Case-to-potential infector distances differed significantly between the observed and permutated data for ESBL-KP, infectors were found more frequently in direct contact (ratio of 3.1, P = 0.009), and no significant difference for ESBL-EC (P = 0.29).

**Table 2.**
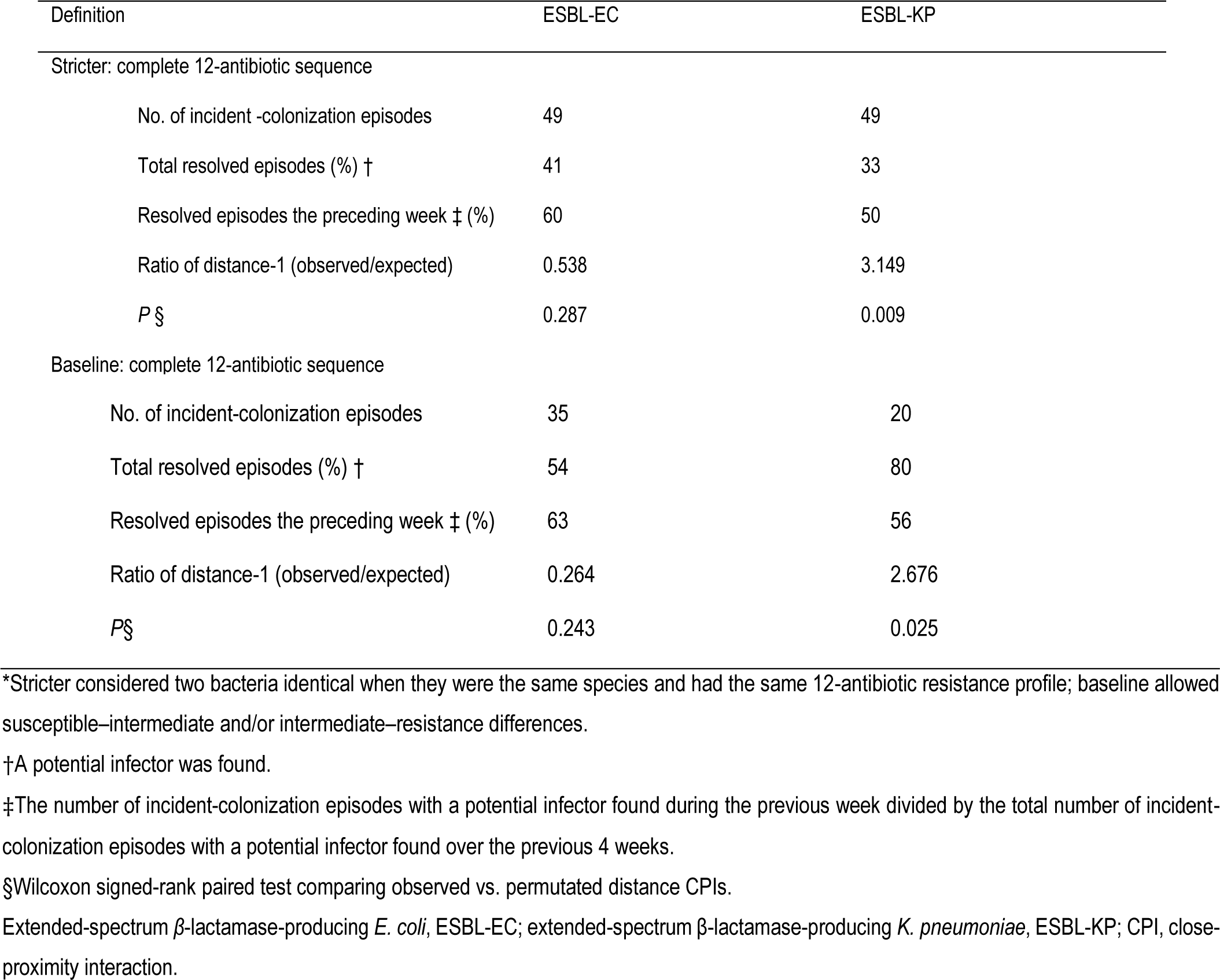
Sensitivity analyses of strain transmission definition according to the stricter or baseline definition^*^.

| Definition | ESBL-EC | ESBL-KP |
| --- | --- | --- |
| Stricter: complete 12-antibiotic sequence |  |  |
| No. of incident -colonization episodes | 49 | 49 |
| Total resolved episodes (%) † | 41 | 33 |
| Resolved episodes the preceding week ‡ (%) | 60 | 50 |
| Ratio of distance-1 (observed/expected) | 0.538 | 3.149 |
| *P* § | 0.287 | 0.009 |
| Baseline: complete 12-antibiotic sequence |  |  |
| No. of incident-colonization episodes | 35 | 20 |
| Total resolved episodes (%) † | 54 | 80 |
| Resolved episodes the preceding week ‡ (%) | 63 | 56 |
| Ratio of distance-1 (observed/expected) | 0.264 | 2.676 |
| *P*§ | 0.243 | 0.025 |

\*Stricter considered two bacteria identical when they were the same species and had the same 12-antibiotic resistance profile; baseline allowed susceptible–intermediate and/or intermediate–resistance differences.
†A potential infector was found.
‡The number of incident-colonization episodes with a potential infector found during the previous week divided by the total number of incident-colonization episodes with a potential infector found over the previous 4 weeks.
§Wilcoxon signed-rank paired test comparing observed vs. permuted distance CPIs.
Extended-spectrum β-lactamase-producing *E. coli*, ESBL-EC; extended-spectrum β-lactamase-producing *K. pneumoniae*, ESBL-KP; CPI, close-proximity interaction.

Sensitivity of the results to the period of investigation. Varying the duration of the investigation period for transmission candidates from 2 up to the entire 25-weeks participation period did not affect the results: more distance-1 than expected by chance were always found for ESBL-KP and never for ESBL-EC (Table 3). The percentages of resolved episodes increased with the investigation-period duration. For instance, for ESBL-KP, 65% of episodes were resolved for the investigation periods was 2-3 weeks, but reached 90% for 8 and 25 weeks.

**Table 3.** Sensitivity analyses of 2-, 3-, 8- or 25-week windows of investigation compared to baseline for transmission candidates.

| Preceding periods | ESBL-EC | ESBL-KP |
| --- | --- | --- |
| 2 weeks |  |  |
| Total episodes resolved*, % | 43 | 65 |
| Resolved episodes found the preceding week†, % | 80 | 69 |
| Ratio of distance-1 (observed/expected) | 0.365 | 2.346 |
| *P*‡ | 0.525 | 0.048 |
| 3 weeks |  |  |
| Total episodes resolved*, % | 51 | 65 |
| Resolved episodes found the preceding week†, % | 67 | 69 |
| Ratio of distance-1 (observed/expected) | 0.292 | 2.905 |
| *P*‡ | 0.468 | 0.057 |
| 4 weeks (baseline) |  |  |
| Total episodes resolved*, % | 54 | 80 |
| Resolved episodes found the preceding week†, % | 63 | 56 |
| Ratio of distance-1 (observed/expected) | 0.264 | 2.676 |
| *P*‡ | 0.243 | 0.025 |
| 8 weeks |  |  |
| Total episodes resolved*, % | 60 | 90 |
| Resolved episodes found the preceding week<sup>†</sup>, % | 57 | 50 |
| Ratio of distance-1 (observed/expected) | 0.420 | 2.179 |
| $P_{\ddagger}^{\dagger}$ | 0.229 | 0.014 |
| 25 weeks |  |  |
| Total episodes resolved*, % | 68 | 90 |
| Resolved episodes found the preceding week<sup>†</sup>, % | 50 | 50 |
| Ratio of distance-1 (observed/expected) | 0.478 | 2.101 |
| $P_{\ddagger}^{\dagger}$ | 0.617 | 0.033 |

\*A potential infector was found.
<sup>†</sup>The number of incident-colonization episodes with a potential infector found during the previous week divided by the total number of incident-colonization episodes with a potential infector found over the 2-, 3-, 4-, 8- or 25-week windows.
<sup>‡</sup>Wilcoxon signed-rank paired test comparing observed vs. expected distances CPIs.
Extended-spectrum β-lactamase-producing *E. coli*, ESBL-EC; extended-spectrum β-lactamase-producing *K. pneumoniae*, ESBL-KP; CPI, close-proximity interaction.

### Simulations of the impact of control measures

We used a mathematical model to assess how control measures would potentially affect our findings. We simulated transmission of an ESBL species in a 100-patient ward over 17 weeks. Without any control measure implemented, the model-predicted cumulative incidence over 4 months was 31% (40/128) patients for ESBL-EC and 19% (24/128 patients) for ESBL-KP, in line with the weekly incidences that were observed during the i-Bird study (Table 2). The four explored illustrative scenarios, based on different levels of isolation and staff hand hygiene, all led to a reduction in incidence. For each control scenario, Figure 2 (Section S7) shows the relative reduction in the 4-month cumulative incidence for both ESBL-EC and ESBL-KP. All scenarios had a significantly larger impact for ESBL-KP than for ESBL-EC, with scenario 3 based on perfect staff hand hygiene being the most effective for both species. Indeed, scenario 3 led to a predicted 39% reduction of the ESBL-KP incidence and a 22% diminution in ESBL-EC incidence, while scenario 1 based on perfect patient isolation led to smaller respective reductions of 14% and 7%. As expected, scenarios 2 and 4 (imperfect compliance) were less effective, with scenario 2 achieving only 12% and 6% reductions of the ESBL-KP and ESBL-EC incidences, respectively.

**Figure 2.**
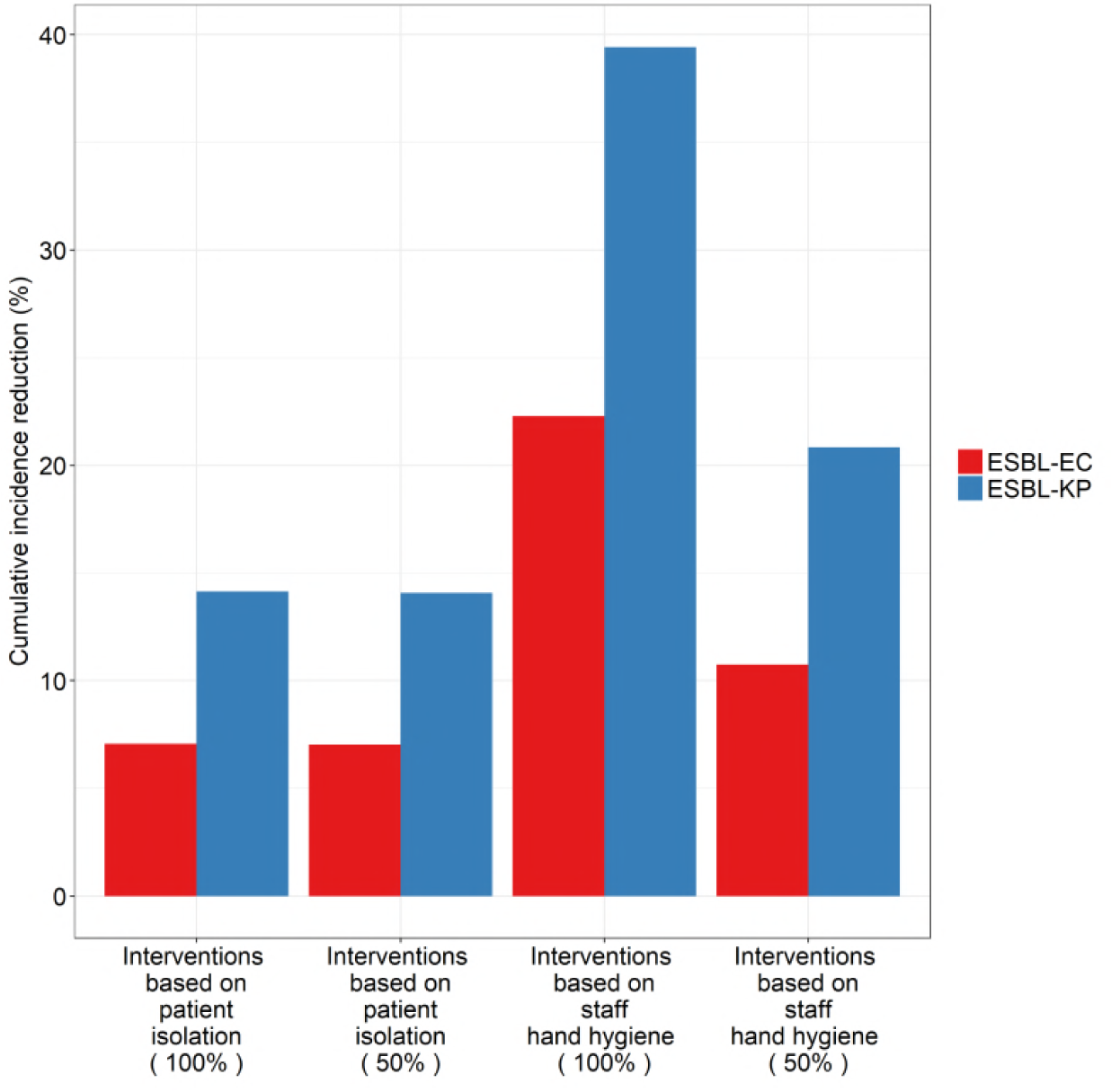
Predicted reduction in the cumulative incidence of ESBL-EC and ESBL-KP under four illustrative scenarios, using the mathematical model. For each scenario and each species, the percent reduction in the cumulative incidence compared to the baseline situation (without any control measure) is depicted (red: EC, blue: KP). Interventions based on patient case isolation correspond to a removal of 100% and 50% of patient-patient CPIs. Interventions based on staff hand hygiene correspond to a removal of 100% and 50% of patient-staff CPIs.

## Discussion

For this study, contact patterns of patients and hospital staff were combined with weekly carriage data to trace the possible routes of resistant Enterobacteriaceae transmission in an LTCF. We found that the human contact network did not correspond to the spread of ESBL-EC, but supported that of ESBL-KP. Those findings suggested that transmission along CPIs is an important driver for ESBL-KP but that it is not the main driver in hospital spread for ESBL-EC. This result is consistent with previous studies investigating the role of patient-to-patient transmission in the Enterobacteriaceae spread (23–25). Because LTCFs can be a hotspot for resistance acquisition (26,27), better understanding of resistant Enterobacteriaceae dissemination in these settings is an important step towards antibiotic-resistance control.

In our study, ESBL-EC and ESBL-KP were the dominant species among ESBL-producing Enterobacteriaceae, in accordance with the reported increase of these two species in Europe over the last years (3). The importation and acquisition rates we observed (8% of admitted patients for importation rate and 0.66%/week acquisition rate for E. coli, 1% importation rate and 0.38%/week acquisition rate for K. pneumoniae) were globally higher than those recently reported in a French intensive care unit for ESBL-producing Enterobacteriaceae as a whole (8% importation rate and 0.29%/week acquisition rate) (28). We also observed a relatively high prevalence of ESBL-EC and ESBL-KP colonization among patients, which is consistent with previous findings in LTCFs (29).

Our results suggest that varied selection and dissemination patterns according to ESBL-producing species. On the one hand, most ESBL-KP acquisitions cases were observed in a specific ward as opposed to broad dissemination of ESBL-EC throughout the entire hospital. On the other hand, the diversity of resistance profiles was broader for ESBL-EC than for ESBL-KP, suggesting potentially higher diversity of circulating E. coli clones, which also agrees with other studies (30). The 8-fold difference between observed E. coli and K. pneumoniae importation rates suggests that the majority of ESBL-EC carriers in our study acquired the strain in the community. This could explain the low portion of resolved ESBL-EC episodes found in our results (9). Another possible explanation for the low ESBL-EC-transmission rate along CPIs is that ESBL-EC might have been acquired mostly through endogenous processes (e.g. plasmid exchange within the gut), after potential resistance acquisition from another species or the environment. Indeed, ESBL-EC is known to represent a resistance-gene reservoir in hospitals (31).

Herein, ESBL-EC and ESBL-KP were analyzed independently, unlike most previous studies in which ESBL-producing Enterobacteriaceae were considered globally, with no species. Although our approach enabled us to highlight the important dissemination differences between the two bacterial species, between-species gene exchanges within a host’s flora were not taken into account, probably contributing to the high unexplained portion of incident colonization episodes with ESBL-EC. Future studies should be designed to specifically assess that question, which will require detailed data on multiple colonization.

This study has several limitations. First, neither the β-lactamase nor its coding gene were identified (or typed), leading us to propose an ad hoc definition of strains based on their phenotypic resistance profiles. As suggested by the sensitivity analyses on strain definition, the impact of that definition on our main results was low. Using genotyping information might not yield different results, especially for E. coli. Indeed, a genetic-based definition of strains would probably be more restrictive and lead to more incident-colonization episodes than resistance phenotypes alone, but with fewer resolved episodes. Hence, our results for ESBL-EC would probably not be affected. For ESBL-KP, because the outbreak remained localized in a specific ward the transmission hypothesis is very likely and despite the genotyping information potentially confirming the outbreak, it is also likely that it would not affect the main study results. Second, CPIs capture all interactions at less than 1.5 m, which means that they do not necessarily involve a physical contact, especially when the CPI duration is short. Thus, it is possible that we captured some false positive “contacts”, especially for patients who shared a room. In general, for most patients, it can be expected that those sharing a room had some contacts with the exception of persistent vegetative state (PVS) patients, who probably had little to no physical contacts with other patients, even while inhabiting the same room. Consequently, transmission between this category of patients might be more likely to occur indirectly, via HCWs or the endogenous route, and captured CPIs between them may not necessarily support bacterial spread. Therefore, it is important to note that a large part of the ESBL-KP–acquisition episodes that were resolved, had a potential infector at a distance-1 for PVS patients (7/13 cases were PVS patients), due to an outbreak of ESBL-KP in neurology ward 1 during to 4-month i-Bird study. More detailed observational data would be needed to fully understand this apparent patient-to-patient transmission of ESBL-KP to PVS patients. In contrast, only 1/35 ESBL-EC– acquisition episodes involved a PVS patient.

Our results have potentially important implications in terms of infection control. Indeed, for ESBL-EC, the most frequently observed case-to-potential infector path was distance-2, with mainly hospital staff intermediaries. Although this might partially reflect the fact that only patients’ swabs were tested for Enterobacteriaceae, the observed pattern of contacts in our LTCF (with frequent patient-patient interactions) suggests that many patient-to-patient CPIs did not result in ESBL-EC transmission. That deduction implies that contact precautions, which are currently the most commonly implemented control measure to prevent ESBL-EC spread (32), may not be fully effective. Our results are consistent with earlier analyses and observations (31,33,34). To further investigate this question, we developed a compartmental model of ESBL-EC or KP spread within a hospital, and simulated two illustrative control measures (patient isolation and staff hand hygiene) with parameters estimated from the study data. Our analyses showed that reducing CPIs, especially between patient and staff (through perfect hand hygiene), might decrease the incidence of both ESBL-EC and ESBL-KP, but with a significantly larger reduction for the latter (Figure 2). Similar results were obtained under various scenarios regarding intervention compliance (isolation with 75%, 50% or 25% of patient-patient contacts removed, hand hygiene with 75%, 50% or 25% of patient-staff contacts removed) (Figure S5). Those findings confirmed that contact-precaution strategies are bound to be highly effective at controlling ESBL-KP, while additional measures such as environmental decontamination or antimicrobial stewardship, might be needed for ESBL-EC.

This study, by jointly analyzing longitudinal ESBL-producing Enterobacteriaceae carriage data with CPI records using radio-frequency identification technology, contributes to our understanding of the dynamics of ESBL-EC and ESBL-KP spread. We showed that CPI information is useful to track ESBL-KP transmission among patients, but not for ESBL-EC. That difference sheds light on the fact that transmission patterns vary according to the species and that species-adapted strategies are necessary needed when aiming to effectively control antibiotic resistance.

## Materials and Methods

### Epidemiological data: The i-Bird study

The i-Bird study was conducted at the Berck-sur-Mer rehabilitation hospital from May 1 to October 25, 2009, with the first 2 months serving as a pilot phase. Rehabilitation centers often require long inpatient periods, unlike acute-care facilities. All participants, 329 patients and 263 hospital staff, wore a badge-sized wireless sensor to record CPIs throughout the study. During that period, rectal swabs were collected weekly from patients to test for Enterobacteriaceae carriage. Hospital staff included all health professionals: healthcare workers (HCWs, including nurses, auxiliary nurses, nurse managers and student nurses), reeducation staff (physical and occupational therapists), ancillary hospital staff, physicians, hospital porters, logistic, administrative and animation staff. The hospital was subdivided into 5 wards, corresponding to medical specialties: neurological rehabilitation (ward 1, 2 and 4), obesity care (ward 3) and geriatric rehabilitation (ward 5).

### CPI description

Every 30 s, each wireless sensor recorded the identification number of all other sensors within a radius of less than 1.5 m and time of interaction. Over 4 months, from July to the end of October, 2,740,728 such distinct CPIs were recorded for 592 persons. This CPI-network was then aggregated at the daily level. To describe CPIs, we used two indicators: the number of daily distinct CPIs of a given individual and the daily cumulative duration of CPI between two individuals. Detailed definitions of these indicators are provided in an earlier paper (35).

### Microbiological data

Rectal swabs were collected weekly from patients. Briefly, swabs were placed in Stuart’s transport medium (500μL; Transwab, Medical 90 Wire and Equipment). Each 100-μL aliquots was plated on selective media for ESBL isolation. The rest of the suspension was then stored at −80°C for further use. Antimicrobial susceptibility testing was done each week for ESBL-producing Enterobacteriaceae, in accordance with national recommendations (36).

### Definitions of carriage

For this study, we independently investigated the spread of two distinct Enterobacteriaceae species: ESBL-EC and ESBL-KP.

#### Strain definition

Strains were defined based on their species characterization and phenotype-resistance profile. Because all strains were ESBL-producing Enterobacteriaceae, we focused on only 12 antibiotics, including 5 aminoglycosides (kanamycin, gentamicin, tobramycin, netilmicin, and amikacin), 4 fluoroquinolones (nalidixic acid, ofloxacin, levofloxacin, and ciprofloxacin), co-trimoxazole, tetracyclin and fosfomycin. Clustering analyses confirmed that the resistance profiles to these 12 antibiotics allowed to define clusters of strains (Section S1 and Figure S2A). In addition, strain profiles were clearly differentiated, with one group dominated by ESBL-EC and the other, more heterogeneous group, had a majority of ESBL-KP (Figure S2B).

Based on those results, we assumed that two ESBL-producing Enterobacteriaceae strains were identical when they belonged to the same species and had the same resistance-sequence status to the 12 selected antibiotics (allowing for R–I or S–I differences for each antibiotic).

#### Definition of prevalence and incidence

Average weekly prevalence and incidence were determined over the 4-month study period for each species and each strain (Section S2). Weekly prevalences were defined as the proportion of colonized patients among swabbed patients during each week of the study period (except for weeks with less than 10 swabbed patient). We defined an incident-colonization episode for a given week as the isolation from a patient of a strain that had not been found in the same patient the preceding week. For a given strain, the weekly incidence was defined as the number of patients with incident-colonization episodes for a given week, divided by the number of patients not colonized by the same strain in the preceding week.

#### Definition of “cases”, “transmission candidates” and “potential infectors”

A “case” was defined as a patient with an incident-colonization episode with a given strain. A “transmission candidate” for a case was defined as a patient who carried the same strain as the case over the preceding 4 weeks (which was the average duration of ESBL carriage in this study). Finally, a case’s “potential infector” was a “transmission candidate” for whom a path linking to the case existed on the CPI-network over the preceding 4 weeks. Thus, potential infectors for a case refer to all individuals who could be at the origin of the strain transmission to the case through the CPI-network. Among all potential infectors, the closest potential infector was the one with the shortest distance to the incident case. When at least one potential infector was found for a given incident case, this case was classified as “resolved”, otherwise it was “unresolved”.

#### Definition of importation and acquisition rates

The importation rate represents the proportion of all admitted patients over the four months of study who were colonized at admission. The weekly acquisition rate is computed as the number of incident colonization episodes among included patients over the four months of study, divided by the total number of included patients over this period and by the study duration, in weeks (Section S3).

### Assessment of the impact of CPIs on transmission of antibiotic resistant bacteria

As described previously (21), the length of the shortest CPI-supported transmission path allows measuring the link between CPIs and bacterial carriage. We tested whether the observed distances between cases and their closest potential infector in the CPI network were different from those expected under the null hypothesis of independence between CPIs and carriage data.

#### Observed distance

For each incident-colonization episode, the observed distance was determined as follows: first, we looked for candidate transmitters carrying the same strain during the preceding weeks. Then, for each candidate, we computed the shortest CPI path (i.e. number of edges) to the case over the last 4 weeks and retained the closest potential infector as the one with the shortest CPI path to compute the distance (Figure 3).

**Figure 3.**
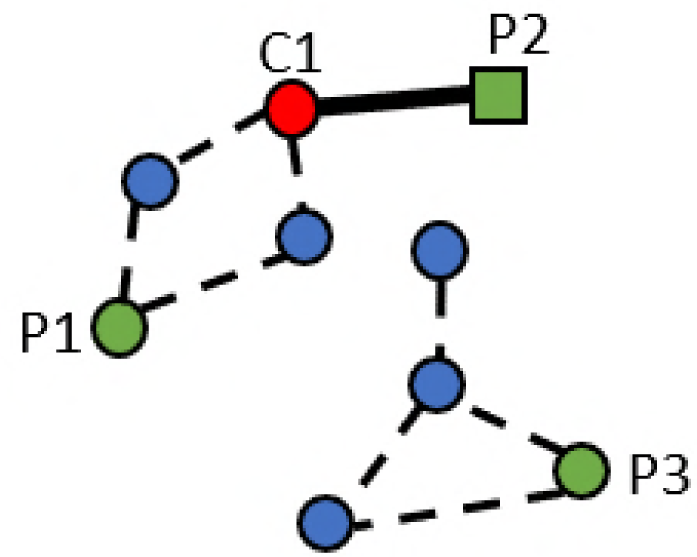
Description of close-proximity interactions (CPIs) and determination of their distances through combined weekly carriage data ans CPI-network plots. Circles and rectangles (nodes) represent patients. The red circle C1 represents a case with an incident - colonization episode. Green circles and rectangle P1, P2, P3 represent transmission candidates, who were colonized with the same strain during the preceding 4 weeks; among them, P1 and P2 represent potential infectors connected to the incident case via edges in the CPI network. Blue circles represent susceptible individuals. Here the closest potential infector is patient P2. The distance is one because no intermediary is present between C1 and P2 (solid black line).

#### Expected distance

For each incident-colonization episode, the expected distance under the null hypothesis was computed through Monte Carlo simulations (Section S4). We randomized all carriage data among the network nodes over the preceding 4 weeks. For each incident-colonization episode, 200 replicates of permutated carriage statuses were obtained. In each permutated dataset, the shortest CPI path was computed as above. The expected distance was then computed by averaging all the shortest lengths of CPI path to this colonization episode.

#### Statistical test

Finally, for all incident-colonization episodes collected, expected distances and observed distances were compared using the Wilcoxon signed rank paired test

### Sensitivity analyses

To assess the influence of assumptions regarding strain definition and colonization duration on the results, CPI-transmission analyses were repeated with different strain definitions and investigation periods. Five outcome indicators were analyzed: (1) the number of incident-colonization episodes; (2) the percentage of resolved incident-colonization episodes (for which potential infectors had been found); (3) the percentage of resolved episodes over the preceding week, which was calculated as the number of incident-colonization episodes for which a potential infector had been found at week 1 divided by the total number of incident-colonization episodes with a potential infector found (Section S5); (4) the ratio of observed versus expected distance-1; and (5) the *P*-value obtained from the Wilcoxon signed rank paired test between observed and expected distances.

First, we compared the five outcome indicators for the results obtained using the initially described strain definition (baseline) in the analysis with those derived with a stricter strain definition. In the later, S–I and I–R differences were taken into account, meaning that for a given incident case, transmission candidates were those carrying strains with the exact same phenotypic resistance profile, as opposed to the less strict baseline definition which allowed those I–S and I–R variations. Then, the impact of the period during which transmission candidates were sought was examined. We repeated the analysis for 2, 3, 4, 8 and the entire 25-weeks study period. The same five indicators were assessed, except for the number of incident-colonization episodes which did not vary according to the considered period duration.

### Deterministic model and simulation

We built a susceptible–colonized model of a 100-patient hospital, in which susceptible (non-colonized) patients could acquire ESBL-producing Enterobacteriaceae following contact with a colonized patient, at a rate *β*_*B*_ for bacteria *B*. *β*_*B*_ was computed as the product of the pathogen-specific per-contact transmission probability (*p*_*B*_) by the weekly distinct number of patients’ CPIs at a distance-1 or distance-2 (*c*_*P*_) observed in the i-Bird CPI-network. Susceptible patients could also become colonized with bacteria *B* at a rate *ν*_*B*_ through the environment or the endogenous route, as previously proposed by Bootsma et al. (37). *ν*_*B*_ was computed as the product of the number of incident colonization episodes for which a potential infector was not found at a distance equal or less than two (1 - *τ*_*B*_), by the weekly incidence rate (*I*_*B*_) of the pathogen observed in the i-Bird data. Colonized patients returned to the susceptible state at a rate *γ*_*B*_, equal in average to 1/*D*_*B*_, where *D*_*B*_ was the duration of bacteria *B* colonization. The model was parameterized for ESBL-EC or ESBL-KP independently. All parameter values were directly taken from the observed i-Bird study data, except for the per-contact transmission probabilities *p*_*B*_, which were computed so that the predicted steady-state colonization prevalence reproduced the observed data (Table 1). More model details, including model equations and details of baseline parameter computation, are provided in Section S6.

We compared the impacts of two simple illustrative control measures with varying levels of compliance, leading to four scenarios: scenarios 1 and 2, in which a portion of patient-patient CPIs was removed to simulate patient contact isolation (scenario 1: 100%, scenario 2: 50%); and scenarios 3 and 4, in which a portion of patient-staff CPIs were removed to simulate staff hand hygiene (scenario 3: 100%; scenario 4: 50%). For each scenario, the mean number *c*_*P*_ of weekly CPIs under a distance-2 was re-computed from the i-Bird data. The corresponding values are provided in Table 4A and Table 4B. All statistical analyses were performed with R version 3.3.2 (http://www.r-project.org/).

**Table 4.**
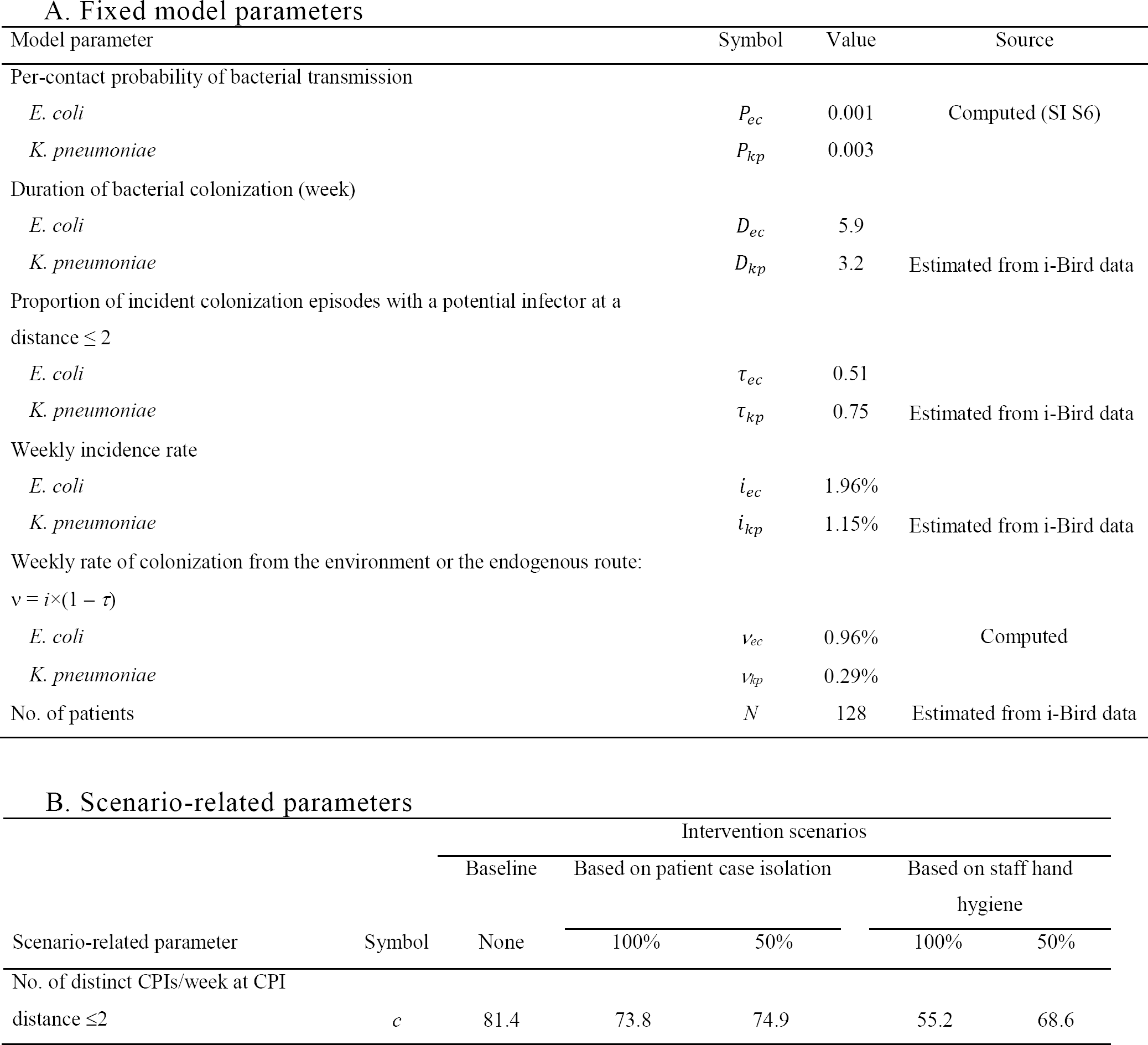
Mathematical model parameters

### Ethics

All authorizations were obtained in accordance with French regulations regarding medical research and information processing. All French IRB-equivalent agencies accorded the i-Bird program official approval (CPP 08061; Afssaps 2008-A01284-51; CCTIRS 08.533; CNIL AT/YPA/SV/SN/GDP/AR091118 N°909036). Signed consent by patients and staff was not required according to the French Ethics Committee to which the project was submitted.

## Supporting Information

Section S1: Resistance profiles of the acquired strains and clustering description Section

S2: Prevalence and incidence definition

Section S3: Importation and weekly acquisition rate

Section S4: Pseudo-code for the calculation of the network-associated distance between an incident colonization episode and potential infector

Section S5: Calculation of the percentage of resolved episodes over the preceding week

Section S6: Mathematical model of bacterial spread within a hospital: model representation, equations, computation of the parameters from the i-Bird data and steady state analysis.

Figure S1 Resistance profile of ESBL-producing Enterobacteriaceae detected in patients over the study period.

Figure S2 - Representation of the model.

Figure S3: Summary of the numbers of colonization, admission, acquisition over the 4-months period.

Figure S4: Number of episodes with a majority of patients or hospital staff intermediaries.

Figure S5: Cumulative incidence reduction from eleven scenarios of the ESBL-EC and ESBL-KP models.

## Acknowledgments

iBird Study Group: Anne Sophie Alvarez (AP-HP, Paris, France), Audrey Baraffe (AP-HP, Paris, France), Mariano Beiró (Universidad de Buenos Aires, Buenos Aires, Argentina), Inga Bertucci (AP-HP, Paris, France), Pierre-Yves Boëlle (Univ. Pierre et Marie Curie, Paris, France), Camille Cyncynatus (AbAg, Chilly-Mazarin, France), Florence Dannet (AP-HP, Paris, France), Marie Laure Delaby (AP-HP, Paris, France), Pierre Denys (AP-HP, Paris, France), Matthieu Domenech de Cellès (Univ. Pierre et Marie Curie, Paris, France), Eric Fleury (ENS Lyon, Lyon, France), Antoine Fraboulet (Insa, Lyon, France), Jean-Louis Gaillard (AbAg, Chilly-Mazarin, France), Boris Labrador (AP-HP, Paris, France), Jennifer Lasley (Inserm, Paris, France), Christine Lawrence (AP-HP, Paris, France), Judith Legrand (Univ. Paris Sud, Orsay, France), Odile Le Minor (Institut Pasteur, Paris, France), Caroline Ligier (Institut Pasteur, Paris, France), Lucie Martinet (Inria, Lyon, France), Karine Mignon (AbAg, Chilly-Mazarin, France), Catherine Sacleux (AP-HP, Paris, France), Jérôme Salomon (Cnam, Paris, France), Thomas Obadia (Univ. Pierre et Marie Curie, Paris, France), Marie Perard (AP-HP, Paris, France), Laure Petit (Institut Pasteur, Paris, France) Laeticia Remy (AP-HP, Paris, France), Anne Thiebaut (Inserm, Paris, France), Damien Thomas (AbAg, Chilly-Mazarin, France), Philippe Tronchet (AP-HP, Paris, France), Isabelle Villain (AP-HP, Paris, France). This study was supported by the European Commission under the Life Science Health Priority of the 6th Framework Program (MOSAR network contract LSHP-CT-2007-037941).

## Author contributions

Conceived and designed the data collection: DG EF PYB. Conceived and designed the analysis: AD, LO, LT. Contributed reagents/analytic tools: JLH. Performed the research: AD. Analyzed the data: AD, TO, LO and LT. Wrote the paper: AD, LO, LT.

## Supporting Information

### Section S1: Resistance profiles of the acquired strains and clustering description

First, the resistance profiles to 28 antibiotics were determined for each identified Enterobacteriaceae from the collected swabs as a sequence of *n* = 28 S, I or R (susceptible, intermediate or resistant, respectively) providing the resistance status to the 28 antibiotics. Because of the wide variability of those profiles among ESBL-producing Enterobacteriaceae and the phenotype detection limits, we clustered bacteria according to their phenotypic profiles. A phenotype distance between two strains was defined by counting the number of R–S mismatches between the strains, ignoring S–I or I–R mismatches.

The distances 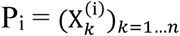 and 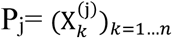 were defined for two resistance profiles with *n* the number of antibiotic resistance phenotypes of the sequence:

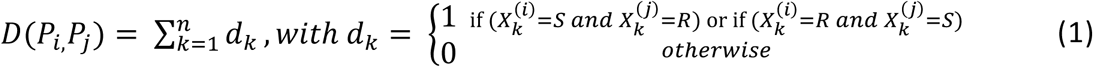

The clustering analysis used the complete linkage method of the function hclust from R version 3.3.2 (http://www.r-project.org/). Strain clusters were based on the calculated distances.

We applied a simplified definition of the sequence and differentiated strain clusters based on a 12-antibiotic resistance profile. That simplified profile includes 5 aminoglycosides (K for kanamycin, GM for gentamicin, TM for tobramycin, NET for netilmicin, and AN for amikacin), 4 fluoroquinolones (NA for nalidixic acid, OFX for ofloxacin, LVX for levofloxacin, and CIP for ciprofloxacin), co-trimoxazole (SXT), tetracycline (TE) and fosfomycin (FOS). With the aim of characterizing strains, bacteria identified as ESBL-EC or ESBL-KP during the study period were also clustered according to this 12-antibiotic–resistance phenotype (SI Appendix, Fig. S1A).

Indeed, in Fig. S1A, different populations may be observed, according to their resistance profiles. In particular, on the upper left part of the figure, strains resistant to aminoglycosides and susceptible to fluoroquinolones on the one hand, and strains susceptible to both antibiotics, appear to form separate groups. In the bottom right side, more heterogeneity is observed. In Fig. S1B, the upper left and bottom right sides can be separated according to the species, with a majority of ESBL-EC on the upper left side and ESBL-KP mostly on the bottom right side. 8 sequences among 96 were shared by ESBL-EC and ESBL-KP

### Section S2: Prevalence and incidence definition

Prevalence and incidence were determined by averaging the weekly values over the *W* weeks of the study period. Let *P* be the total number of patients included in the study. For each week *w* (in 1…*W*) and any patient *p* (in 1…*P*), let *P*_*wp*_ be an indicator of presence within the hospital of patient *p* during week *w* (*P*_*wp*_ = 1 if patient *p* was present), and let *C*_*wp*_ be an indicator of colonization for patient *p* on week *w* (*C*_*w p*_ = 1 if patient *p* was colonized). Then the weekly prevalence and incidence during week *w* (in 1…*W*) can be computed as:

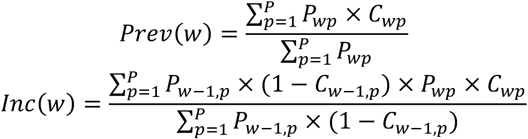

The average weekly prevalence and incidence over the study period can be computed as:

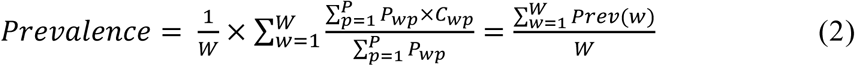

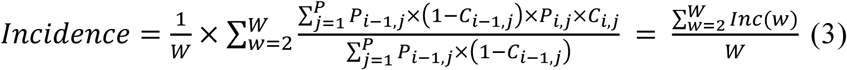

### Section S3: Importation and weekly acquisition rate

Global importation and acquisition rates were calculated over the *W* weeks of the study period as follows:

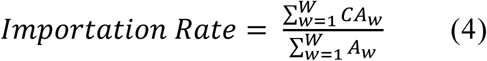

where *CA*_*w*_ is the number of colonized participating patients admitted during week *w*, and *A*_*w*_ is the number of participating patients admitted during week *w*.

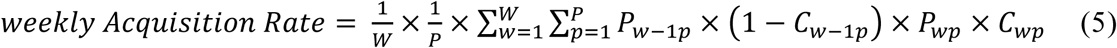

where *P* is the total number of patients included in the study, and for any week *wp* (in 1…*W*) and any patient *p* (in 1…*P*), *P*_*wp*_ is an indicator of presence within the hospital of patient *p* during week *w* (*P*_*wp*_ =1 if patient *p* was present), and *C*_*wp*_ is an indicator of colonization for patient *p* during week *w* (*C*_*wp*_ =1 if patient *p* was colonized).

### Section S4: Pseudo-code for the calculation of the network-associated distance between an incident colonization episode and potential infector For each incident colonization episode *i* = 1 to Ncases do

0. *Identify the episode*

Let the concerned patient as *p*^*i*^, the date as *w*^*i*^ and the microbiological result as *m*^*i*^

*1.Find all swabs from individuals other than* p^i^ *taken during the time window [*w^i^*–*W, w^i^*–*1] Tab = table of patient IDs/swab dates/microbiological results

#### # To compute the observed distance

*2. Find all transmission candidates in Tab*

For each swab in Tab

If microbiological result (swab) = *m*^*i*^Add swab to CandidateTab

*3.Compute the distance* dt *from the episode to each transmission candidate* t

For each transmission candidate *t* in Candidate Tab

*dt* = 1

While [(found = FALSE) and (*dt* < DMAX)] repeat IDlist = list of all patient IDs with network links of length *dt* to patient *p*^*i*^ in the time window [*w*^*i*^–*W, w*^*i*^–1]

If *t* is in IDlist do Set found = TRUE Else do*dt* = *dt* + 1

*4. Determine the distance to the closest potential infector*

Compute *d* as the minimum of all *dt*

#### # To compute the expected distance under H0 Do *n* times

*5.Shuffle the microbial data*

Randomly shuffle the last column of Tab

*6.Perform steps 2 to 4 to compute the episode’s distance to its closest potential infector* 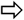 **Compute *d*_*exp*_ as mean of the *n* distances**

### Section S5: Calculation of the percentage of resolved episodes over the preceding week

The percentage of resolved episodes during the preceding week represents the proportion of episodes with a potential infector found during the preceding week among all incident-colonization episodes that were detected during the entire study detection period. We calculated this percentage as follows

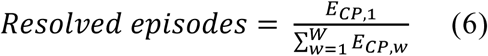

Where *W* is the total number of weeks in the study, *E*_*CP,1*_ is the number of incident-colonization episodes with a potential infector found during the preceding week and *E*_*CP,i*_is the number of incident-colonization episodes with a potential infector found during week *w*.

### Section S6: Mathematical model of bacterial spread within a hospital: model representation, equations, computation of the parameters from the i-Bird data and steady state analysis

#### Notations

Let *C(t)* be the number of patients within the hospital colonized with bacteria *B* at time *t*, and *S(t)* the number of patients not colonized with that bacterium, and therefore susceptible to acquire its colonization. At all times: *S(t)* +*C(t)* = *N*, the total number of patients within the hospital. Let *β*_*B*_ be the weekly effective contact rate, computed as:

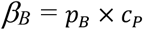

where *c*_*P*_ is the per-patient average weekly number of CPIs at a distance ≤ 2, and *P*_*B*_is the per-contact transmission probability of bacteria *B*.

Let *v*_*B*_ be the weekly colonization-acquisition rate via the endogenous route or the environment, computed as:

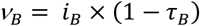

where *i*_*B*_ is the observed weekly incidence rate of bacteria *B* and τ_*B*_ is the proportion of cases of incident-colonization with bacteria *B* for which a potential infector was found at a distance ≤ 2. Let *γ*_*B*_be the decolonization rate of bacteria *B*, computed as:

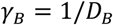

where *D*_*B*_ is the average duration (in weeks) of colonization with bacteria *B*, estimated as the ratio of the average prevalence of colonization with bacteria *B* by the average weekly incidence of colonization with bacteria *B*.

#### Model equations

Time changes in *S(t)* and *C(t)* are driven by the following differential equations:

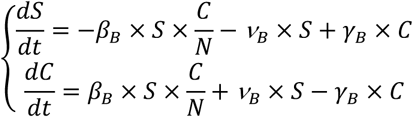

#### A schematic representation of the model is presented on Fig S2

<u>***Computation of P*_*B*_**</u>

The steady-state values of *S* and *C*, denoted as *S*^*\**^ and *C*^*\**^, verify the following equation:

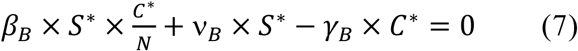

Let *s*^*\**^ and *c*^*\**^ be the steady-state proportions of susceptible and colonized patients within the hospital:

*s*^*\**^ *= S*^*\**^*/N* and *c*^*\**^ *= C*^*\**^*/N*.

Then it can be inferred from (7) that:

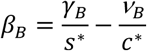

And as *β*_*B*_ = *P*_*B*_ *× c*_*P*_, the per-contact probability of bacteria *B* transmission *p*_*B*_ can be computed as:

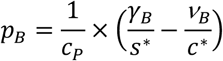

#### Numerical application for ESBL-EC

Based on the i-Bird CPI data, the baseline per-patient average weekly number of CPIs at a distance ≤ 2 is *c*_*P*_ = 81.4 CPIs/week.

The average prevalence and weekly incidence of ESBL-EC in the i-Bird study, considered as steady-state values, are (from Table 1):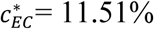 and *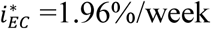*

Hence, the duration of ESBL-EC colonization may be estimated as:

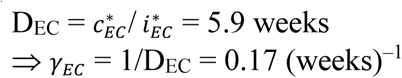

Based on our analysis of incident cases and potential infectors, the proportion of cases of incident ESBL-EC colonization with a potential infector identified at a distance ≤ 2 is:

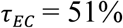

Hence the weekly rate of ESBL-EC acquisition from the endogenous route is estimated at:

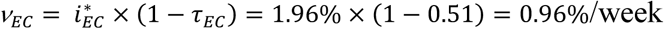

And the per-contact probability of ESBL-EC transmission may be computed as:

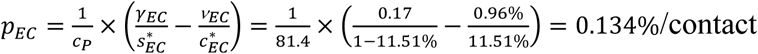

#### Numerical application for ESBL-KP

Based on the i-Bird CPI data, the baseline per-patient average weekly number of CPIs at a distance ≤ 2 is *c*_*P*_=81.4 CPIs/week.

The average prevalence and weekly incidence of ESBL-KP in the i-Bird study, considered as steady-state values, are (from Table 1):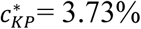 and *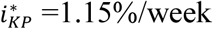*

Hence, the duration of ESBL-KP colonization may be estimated as:

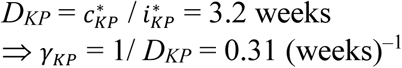

Based on our analysis of incident cases and potential infectors, the proportion of cases of incident ESBL-KP colonization with a potential infector identified at a distance ≤ 2 is:

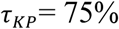

Hence the weekly rate of ESBL-KP acquisition from the endogenous route or the environment is estimated at:

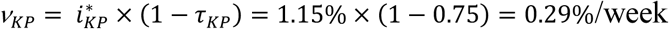

And the per-contact probability of ESBL-KP transmission may be computed as:

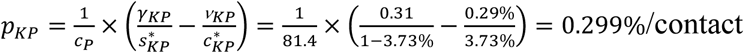

### Model Output: Reduction in the cumulative incidence

For each scenario *s* and each bacterium, the model was run for 17 weeks and the cumulative incidence of acquisitions was calculated 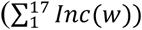 Then the reduction of cumulative incidence for a given scenario compared with a scenario with no intervention was calculated as follows:

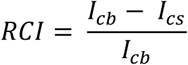

Where *I*_*cb*_ represents the 4-month cumulative incidence under the baseline scenario and *I*_*cs*_ the 4-month cumulative incidence under the intervention scenarios.

**Fig. S1.**
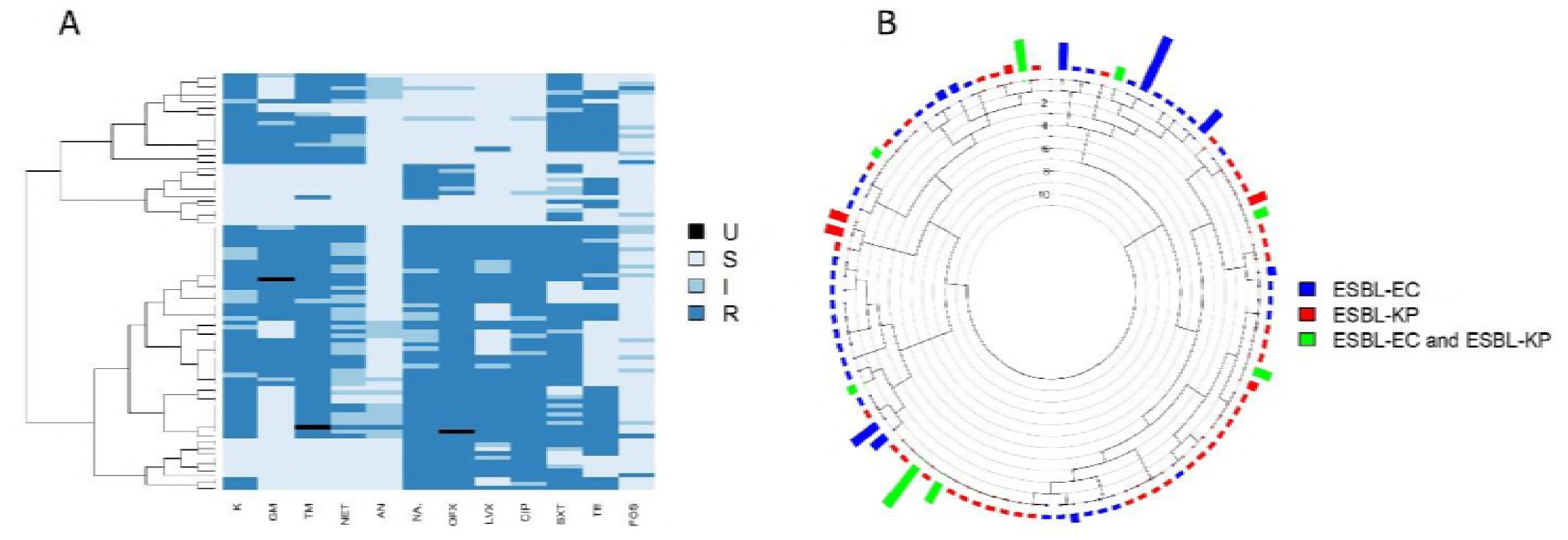
Resistance profile of ESBL-producing Enterobacteriaceae detected in patients over the study period. (A), Each row represents a strain identified during the study. Each columns represents the phenotype sequence in terms of antibiotic resistance level to each of the 12 tested antibiotics. R, resistant (dark blue). I, intermediate (blue). S, susceptible (light blue) and U unknown (black). Tested antibiotics were penicillins (aminoglycosides (kanamycin (K), gentamicin (GM), tobramycin (TM), netilmicin (NET), amikacin (AN)), fluoroquinolones (nalidixic acid (NA), ofloxacin (OFX), levofloxacin (LVX), ciprofloxacin (CIP)), co-trimoxazole (SXT), tetracyclines (TE) and fosfomycin (FOS). The dendrogram was built from the distances between two strains’ phenotype profiles for the 12 antibiotics. (B) The same data is represented with characterization of the species. Blue: ESBL-EC, red: ESBL-KP and green: resistance sequences found in both species. Rectangle heights correspond to the number of individuals each profile was observed in.

**Fig. S2.**
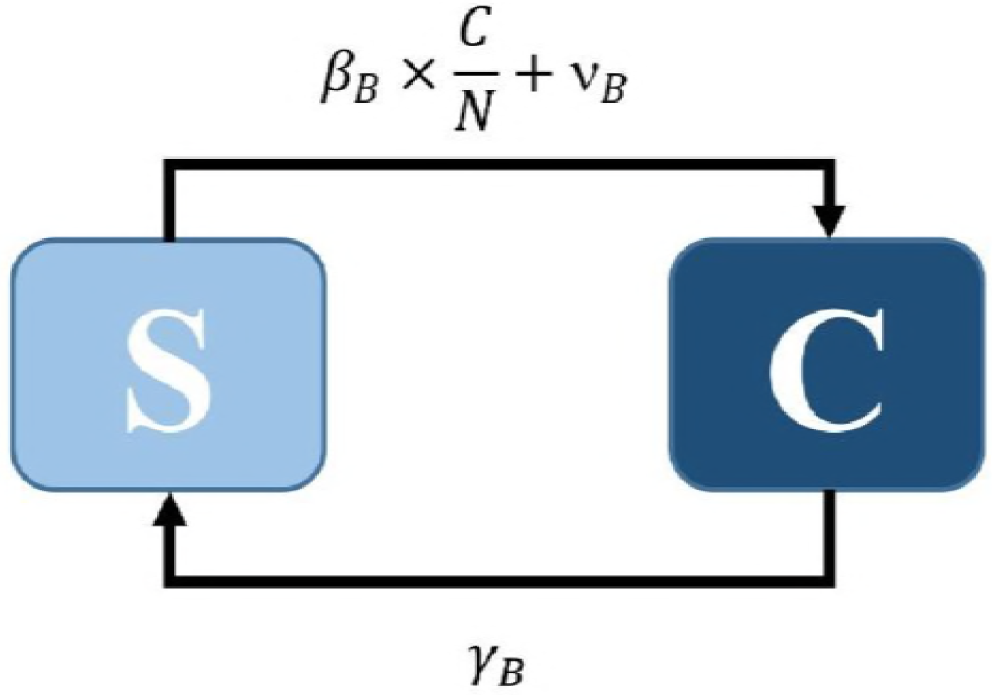
Representation of the model. S and C are the susceptible and colonized compartments. β_B_ is the weekly effective contact rate, N is the total number of patients within the hospital, *v*_*B*_ is the weekly colonization-acquisition rate via the endogenous route or the environment and γ_*B*_ is the decolonization rate of bacteria B.

**Fig. S3:**
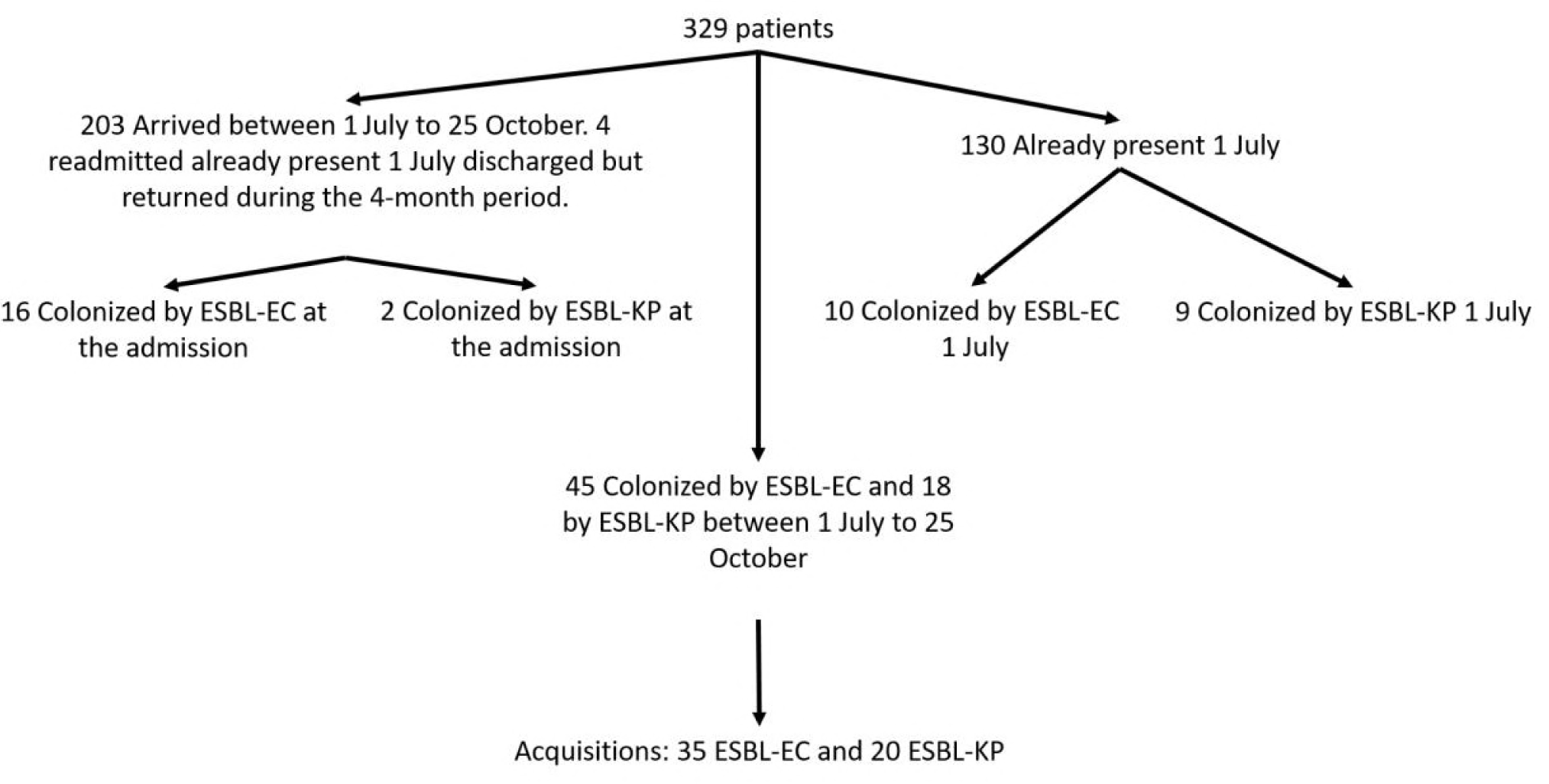
Summary of the numbers of colonization, admission, acquisition over the 4-months period.

**Fig. S4:**
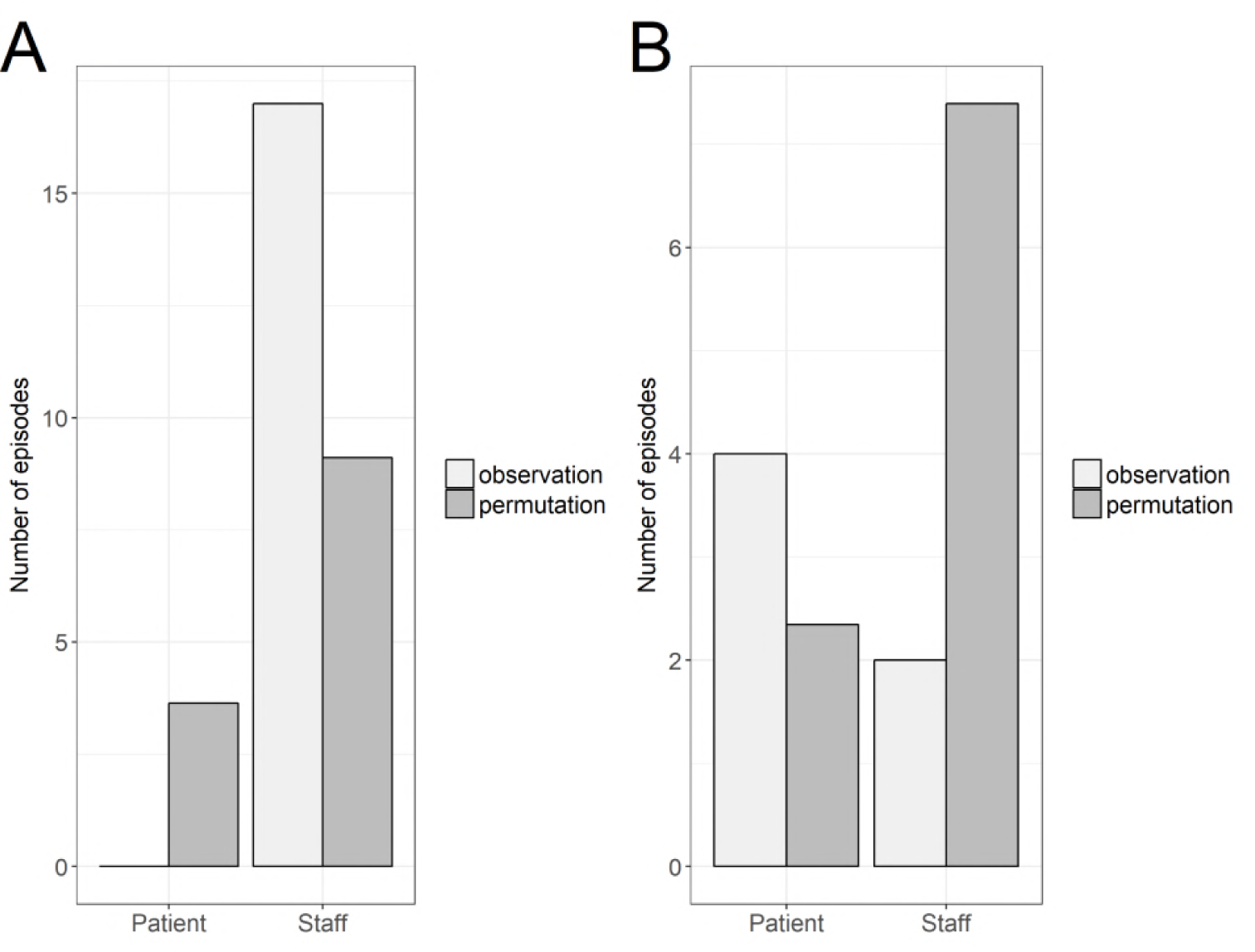
Number of episodes with a majority of patients or hospital staff intermediaries. Here, only incident colonization episodes with a distance-2 to their potential infector are considered. The portions of these episodes in which there is a majority of patients and hospital staff are depicted for (A) ESBL-EC and (B) ESBL-KP. These portions are compared between observed (light gray) and randomly permutated data (dark grey).

**Fig. S5:**
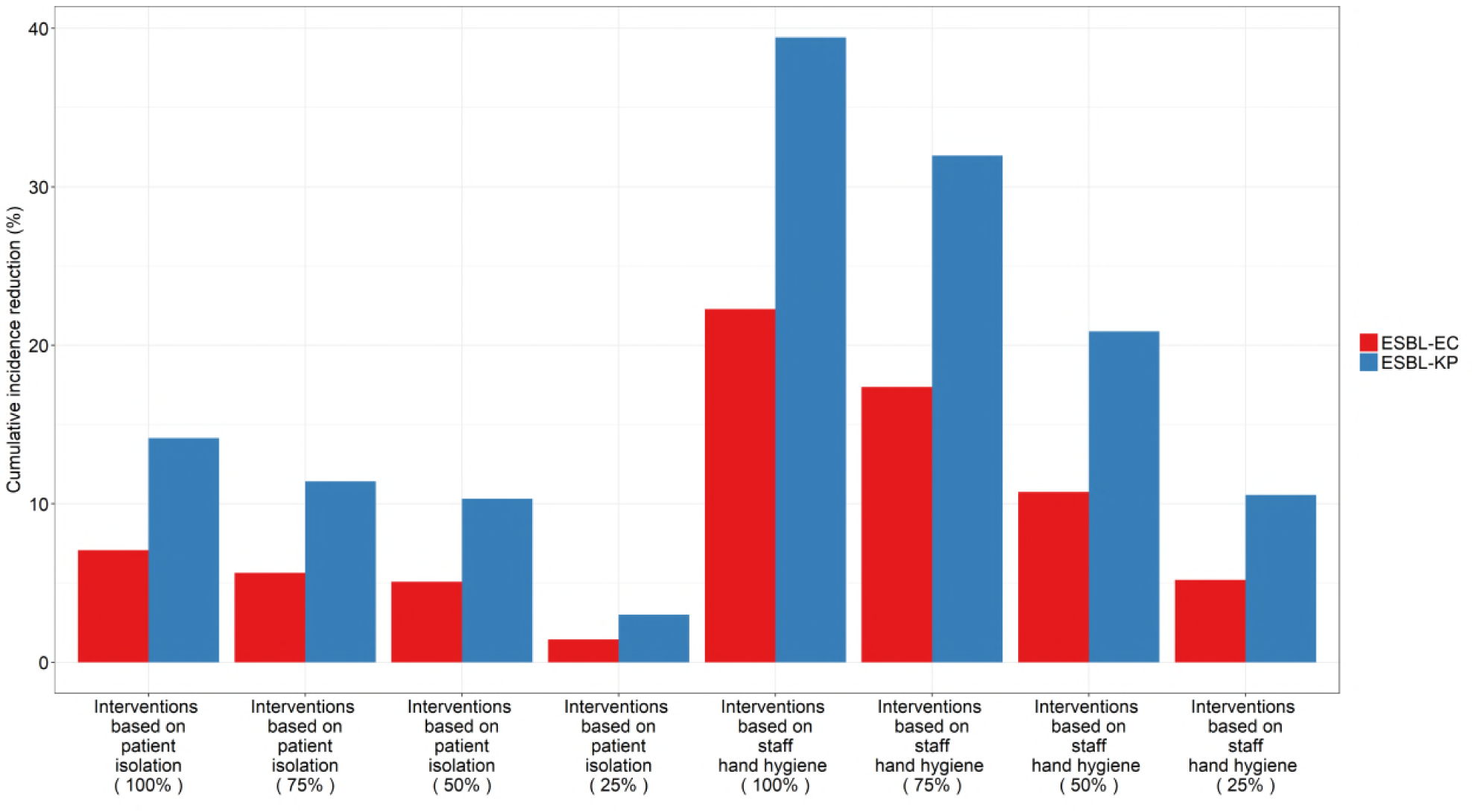
Cumulative incidence reduction from eleven scenarios of the ESBL-EC and ESBL-KP models. Percentage on the y-axis corresponds to reduction of the cumulative incidence compared to the baseline (scenario with no control measure). In red, percentage of cumulative incidence reduction of ESBL-EC and blue ESBL-KP. Intervention based on patient isolation correspond to a removal of 100%, 75%, 50% and 25% of patient-patient CPIs. Intervention based on staff hand hygiene correspond to a removal of 100%, 75%, 50% and 25% of patient-staff CPIs.

